# A novel role of MNT as a negative regulator of REL and the NF-κB pathway

**DOI:** 10.1101/2020.07.21.210989

**Authors:** Judit Liaño-Pons, M. Carmen Lafita-Navarro, Carlota Colomer, Lorena García-Gaipo, Javier Rodríguez, Alex von Kriegsheim, Peter Hurlin, M. Dolores Delgado, Anna Bigas, M. Lluis Espinosa, Javier Leon

## Abstract

MNT, a transcription factor of the MXD family, is an important modulator of the oncoprotein MYC. Both MNT and MYC are basic-helix-loop-helix proteins that heterodimerize with MAX in a mutually exclusive manner, and bind to E-boxes within regulatory regions of their target genes. While MYC generally activates transcription, MNT represses it. However, the molecular interactions involving MNT as a transcriptional regulator beyond the binding to MAX remain unexplored. Here we demonstrate a novel MAX-independent protein interaction between MNT and c-REL (REL), the oncogenic member of the REL/NF-κB family. REL is involved in important biological processes and it is found altered in a variety of tumors. REL is a transcription factor that remains inactive in the cytoplasm in an inhibitory complex with IκB and translocates to the nucleus when the NF-κB pathway is activated. In the present manuscript, we show that *MNT* knockdown triggers REL translocation into the nucleus and thus the activation of the NF-κB pathway. Meanwhile, *MNT* overexpression results in the repression of IκBα, a *bona-fide* REL target. Indeed, both MNT and REL bind to the IκBα gene at a region mapping in the first exon, suggesting its regulation as a MNT-REL complex. Altogether our data indicate that MNT acts as a repressor of the NF-κB pathway by two different mechanisms: 1) retention of REL in the cytoplasm by MNT protein interaction and 2) MNT-driven repression of REL-target genes through a MNT-REL complex. These results widen our knowledge about MNT biological roles and reveal a novel connection between the MYC/MXD and the NF-κB pathways, two of the most prominent pathways involved in cancer.

## INTRODUCTION

MNT is a protein from the MYC/MAX/MXD/MLX network of transcription factors, which plays a pivotal role in controlling cell proliferation, differentiation, metabolism and oncogenic transformation. MNT is a basic helix-loop-helix leucine zipper (bHLHLZ) protein that regulates transcription as homodimers^1^ and heterodimers with MAX ^2,3^ or MLX^4^. Hence, MNT connects the MYC-MAX and MLX-Mondo branches of the network^5^. MNT normally represses gene transcription by binding to E-boxes and interacting with SIN3 proteins that, in turn, recruit histone deacetylase complexes to its target genes^6,7^.

Among the MXD proteins, MNT is the biggest as well as the most ubiquitously expressed and conserved member^8,9^. Whereas *Mxd1*^−/−^, *Mxi1*^−/−^ and *Mxd3*^−/−^ mice survive, mice knockout for Mnt die soon after birth^10-13^. Thus, MNT is a unique and essential protein of this network. MNT is also frequently deleted in cancer, e.g. in chronic lymphocytic leukemia, Sézary Syndrome (a variant of cutaneous T-cell lymphoma) and medulloblastoma^14-17^. Indeed, around 10 % of the tumors show deletions of a MNT allele^18^.

MNT plays an important role in modulating the oncogenic activities of MYC whether as an antagonist and tumor suppressor or as a cooperator^8^. MNT-MYC antagonism is achieved at three different levels: (i) competition for binding to MAX; (ii) competition between MNT-MAX and MYC-MAX for binding to the E-Boxes of their shared target genes; (iii) transcriptional repression of shared target genes that are normally activated by MYC-MAX^10,11^. This antagonism can explain why the deletion of MNT leads to tumor formation in mouse mammary epithelium and T-cells^10,11^. However, other studies suggest that MYC needs the pro-survival functions of MNT for fully achieving its transformation potential. This is the case of MYC-driven B- and T-cell lymphoma models, where MNT deficiency impairs MYC-driven tumorigenesis^10,11,19^

Nevertheless, there are several unsolved questions about MNT mechanism of action. All the functions described so far for MNT have been attributed to MNT-MAX dimers. However, MAX is deleted in some cancers, as pheochromocytoma, paraganglioma, gastrointestinal stromal tumors and small cell lung cancer^20,21^. Moreover, we have recently described MAX-independent MNT activities in cell proliferation and gene transcription^1^. Thus, we hypothesized that there are MNT functions dependent on the interaction with other proteins different from MAX. In this work we have investigated new MNT interactions in a MAX-independent setting and identified c-REL (REL hereafter), a member of the NF-κB’s family, as a MNT interacting protein. NF-κB signaling pathway plays a major role in proliferation, differentiation and apoptosis particularly in cells from the immune system^22,23^. REL was first described by homology with v-*rel*, the oncogene from the avian reticuloendotheliosis virus^24^. Indeed, REL is the only NF-κB protein with transforming ability^25^ and is altered in a variety of tumors ^26^. We also show that MNT deletion leads to the translocation of REL into the nucleus and to the activation of the NF-κB pathway. Moreover, MNT provokes the repression of NF-κB signaling and the binding, together with REL, to NF-κB target gene, *NFKBIA*/IκBα, a protein required to retain NF-κB dimers in the cytoplasm in the absence of activating stimulus^23^. In summary this work describes the first evidence of a physical interaction between MYC-MAX-MNT and the NF-κB pathways and provides an insight into the relevance of MNT in cell biology.

## RESULTS

### Searching for new MNT partners

To find proteins that interact with MNT in a MAX-independent way, we performed proteomic analysis of MNT immunoprecipitates in UR61-derived cells, which come from a rat pheochromocytoma and lack a functional MAX gene^27^. We used URMax34, a cell line with a Zn^2+^-inducible MAX allele and its control cells (URMT), which were transfected with the empty vector^1^. MNT was immunoprecipitated by triplicate for each cell line and subjected to mass spectrometry. The results showed 11 proteins that were reproducibly immunoprecipitated in URMT and 47 in URMax34 (Supplementary Table 1). Five proteins were shared between the two cell lines: REL, CCDC6, AMPD2, QSER1 and TPP2 (Fig. 1a). Given the relevance of the NF-κB pathway in cell biology, we selected REL for further studies. First, we tested the protein expression of MNT and REL levels in several cell lines, including URMT and URMax34 cells, by western blot (Fig. 1b). As we recently described, MNT expression was higher in the URMT cells lacking MAX ^1^. Next, we analyzed the oncoprint of MNT and REL with the cBioPortal tool ^28^. Mutations in these genes were found in 320 samples from the Cancer Genome Atlas Program (TCGA) database and they were mostly mutually exclusive (Fig. 1c), suggesting that both genes could act in the same pathway. Moreover, we observed a negative correlation between the impact of MNT and REL levels on the survival in some tumor types (Supplementary Fig. S1).

**Figure 1.**
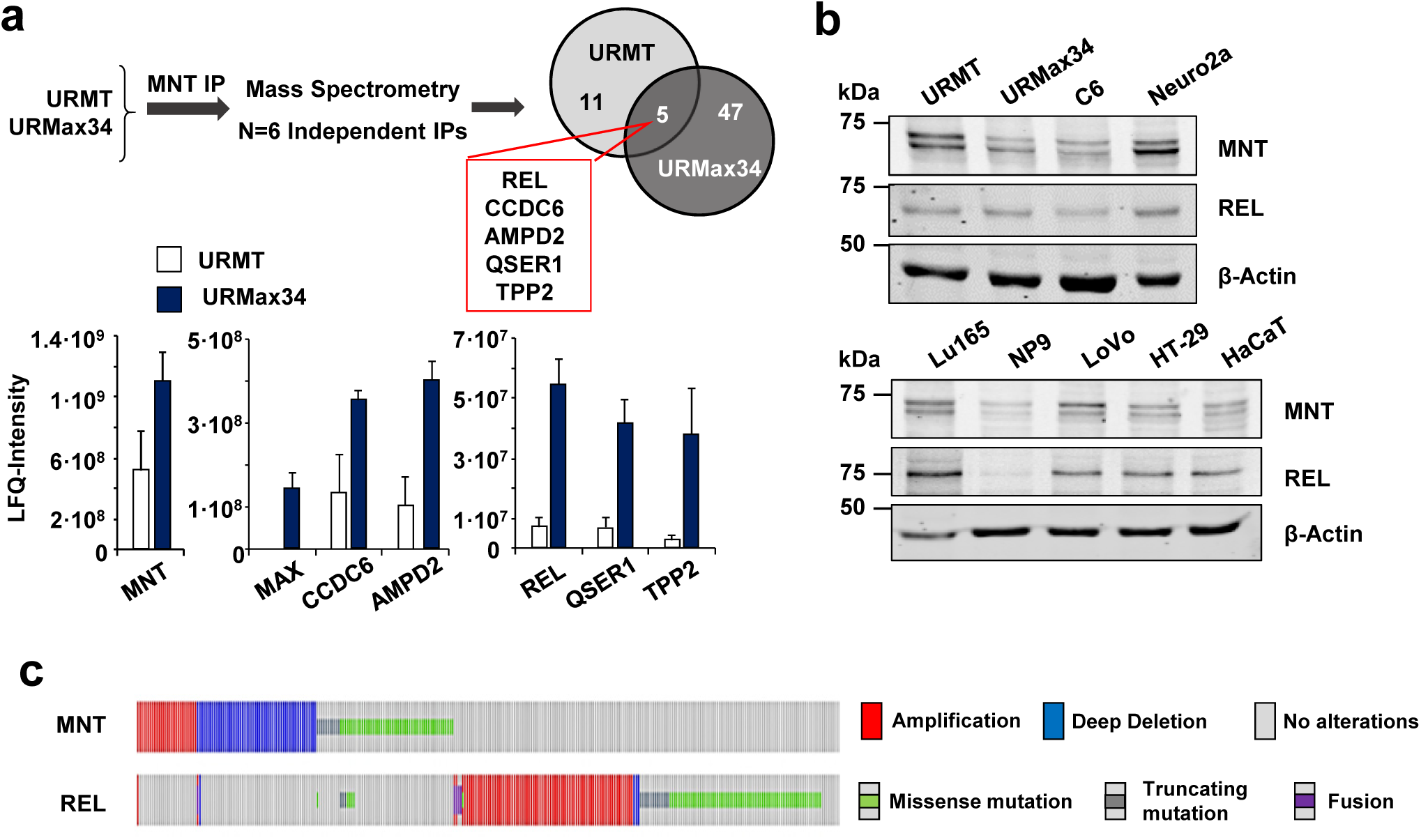
Proteins interacting with MNT. **(a)** Top: Schematic representation of the proteomic study: URMT and URMax34 cells were treated with 100 µM Zn_2_SO_4_ for 24 h and lysed for posterior MNT immunoprecipitation. Six independent immunoprecipitations were analysed by mass spectrophotometry and 5 proteins were found bound to MNT in URMT and URMax34. Bottom: Quantification of the protein interactions. The bar graphs represents mean LFQ intensity values of the selected Mnt interactions ±S.D. (n = 6). P < 0.05 with respect to IgG. **(b)** Immunoblot of MNT and REL in different cell lines: URMT (control UR61) and URMax34 (UR61 with MAX expression induced by Zn^2+^); C6 (rat brain glioma); Neuro-2a (mouse neuroblastoma); Lu165 (small cell lung cancer); NP9 (human pancreatic adenocarcinoma); LoVo and HT-29 (human colorectal adenocarcinoma); HaCaT (human keratinocyte). β-actin levels were determined as protein loading control. **(c)** Oncoprint for MNT and REL obtained from cBioPortal (cbioportal.org). The individual genes are represented as rows, and individual cases (320 in total, TCGA, PanCancer Atlas) are represented as columns.

To confirm the MNT-REL interaction, we performed co-immunoprecipitation (co-IP) assays followed by western blot. REL was present in MNT immunoprecipitates of URMax34 cells induced to express MAX. In this model we confirmed the MNT-MAX co-IP. However, REL was not present in the MAX immunoprecipitates, indicating that there was no interaction of REL with MAX (Fig. 2a). We also found REL-MNT co-IP in mouse Neuro-2a (Fig. 2b), human colon cancer LoVo (Fig. 2c) and rat glioma C6 cells (Fig. 2d, left). MNT-REL interaction was not detected in other cell lines tested, e.g., SH-SY5Y (human; neuroblastoma), HEK293T (human; embryonic kidney), K562 (human; chronic myelogenous leukaemia), Jurkat (human; acute T cell leukemia), HaCaT (human keratinocytes), A549 and H1299 (human; NSCLC), H1417, LU165 (human; SCLC), HEPG2 (human; hepatocellular carcinoma), Rat1a (rat fibroblasts), MEFs (mouse embryonic fibroblasts), COS7 (monkey; kidney fibroblasts).

**Figure 2.**
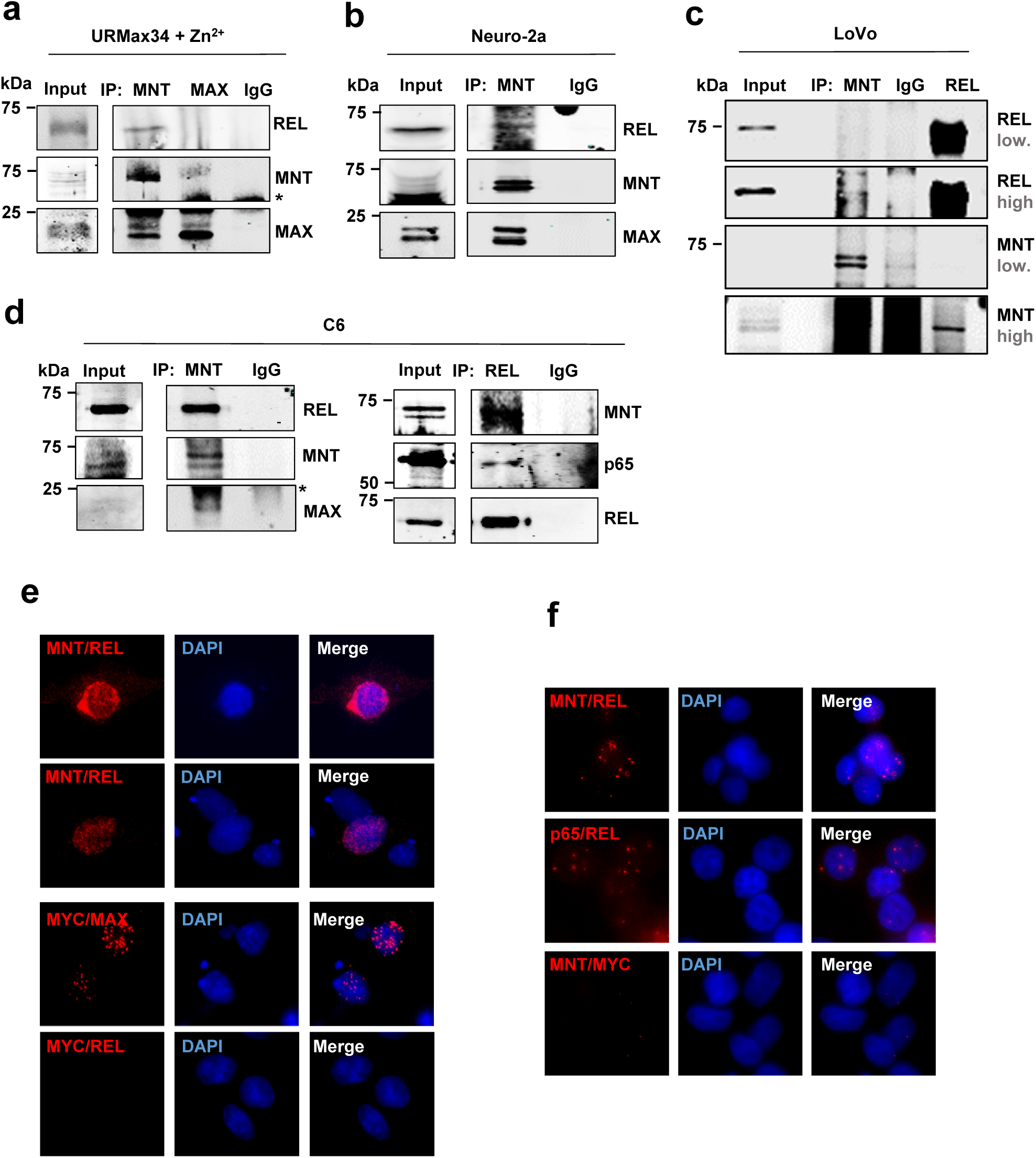
MNT and REL physically interact. **(a)** Co-immunoprecipitation assay with anti-MNT and anti-MAX antibodies in URMax34 cells after Zn2+ treatment. MAX was used as a positive control for MNT immunoprecipitations. The asterisk marks the position of the heavy IgG band. **(b)** Co-immunoprecipitation assay with anti-MNT in mouse Neuro-2a cells. **(c)** Co-immunoprecipitation assay in human LoVo cells with anti-MNT and antgi-REL. Two immunoblot images of the blots with high and low intensity are shown. Note that the hypotonic buffer required for MNT-REL co-immunoprecipitation does not extract efficiently MNT protein. **(d)** Co-immunoprecipitation assay in rat glioma C6 cells with anti-MNT and anti-REL antibodies **(e)** Proximity Ligation assay performed in C6 cells 48 h after transfection with WT MNT-HA and REL-flag overexpressing vectors. Antibodies anti-MNT/anti-REL, anti-MYC/anti-MAX (positive control) and anti-MYC/anti-REL (negative control). PLA positive signal in red and DAPI as a nuclear marker. **(f)** Proximity Ligation Assay in LoVo cells (untransfected cells) with anti-MNT/anti-REL, anti-p65/anti-REL (positive control) and anti-MNT/anti-MYC (negative control) antibodies. PLA positive signal in red and DAPI as a nuclear marker.

We also detected MNT-REL interaction in immunoprecipitates with anti-REL antibodies in C6 (Fig. 2d, right) and URMT cells (not shown). Next, we wanted to confirm the interaction between MNT and REL through Proximity Ligation Assays. We transfected C6 cells with MNT and REL expression vectors and obtained a positive result of MNT-REL interaction (Fig. 2e). A positive result was also obtained in untransfected LoVo cells, although to a lesser extent (Fig. 2f). The interactions between MYC-MAX and p65-REL were used as positive controls of the assay. MYC-REL and MNT-MYC were used as negative controls.

REL is generally found forming homodimers or heterodimers with p65 or p50 ^22^. To assess if there was any p65 or p50 in the MNT-REL complex, we immunoprecipitated p65 to show the p65-REL interaction in UR61 cells with and without MAX. We observed p65-REL interaction but not the p65-MNT co-IP (Fig. 3a). We also performed co-IP assays using two different antibodies against MNT, recognizing the first 1-50 amino acids and the 532-582, respectively, in LoVo cells. In addition, we used antibodies anti-p65 and p105/p50. The results showed the REL-MNT co-IP with the anti-MNT antibody recognizing the MNT 1-50 amino acids but not in the anti-MNT recognizing the amino acids 532-582. The binding of the antibody to the 532-582 amino acids of MNT may disrupt MNT-REL co-IP, which suggests the implication of MNT C-terminal domain in the interaction. MAX co-IP was detected in both MNT immunoprecipitates (positive control) (Fig. 3b). However, neither p65 nor p50 were found to interact with MNT in any of the IPs. REL co-immunoprecipitated with p65 and p50 as expected. Since p65 and p50 did not interact with MNT, it is possible that the MNT-REL complex may be composed by REL homodimers or REL bound to other unknown protein(s), as schematically represented in Fig. 3c. To determine the domain of MNT involved in the interaction with REL, we used two MNT deletion mutants tagged with the HA epitope: ΔbHLH (lacking the bHLH domain, residues 221-272) and ΔCt1 MNT-HA (lacking the C-terminal region of 276 residues) (Fig. 3d). We transfected these constructs along with a construct expressing mouse REL into C6 cells. Interestingly, REL appeared bound to ΔbHLH but not to ΔCt1 (Fig. 3e). Altogether, these data show that the C-terminal region of MNT is necessary for the interaction. Next, we investigated the localization of MNT-REL complexes carrying out a nucleus-cytoplasm fractionation in C6 cells. The co-IP was performed in basal conditions and after 30 min of stimulation with TNFα (which promotes the translocation of REL to the nucleus)^29^. The results showed that the MNT-REL complex localizes in the cytoplasm in normal conditions but also in the nucleus if the NF-κB pathway is activated by TNFα (Fig. 3f).

**Figure 3.**
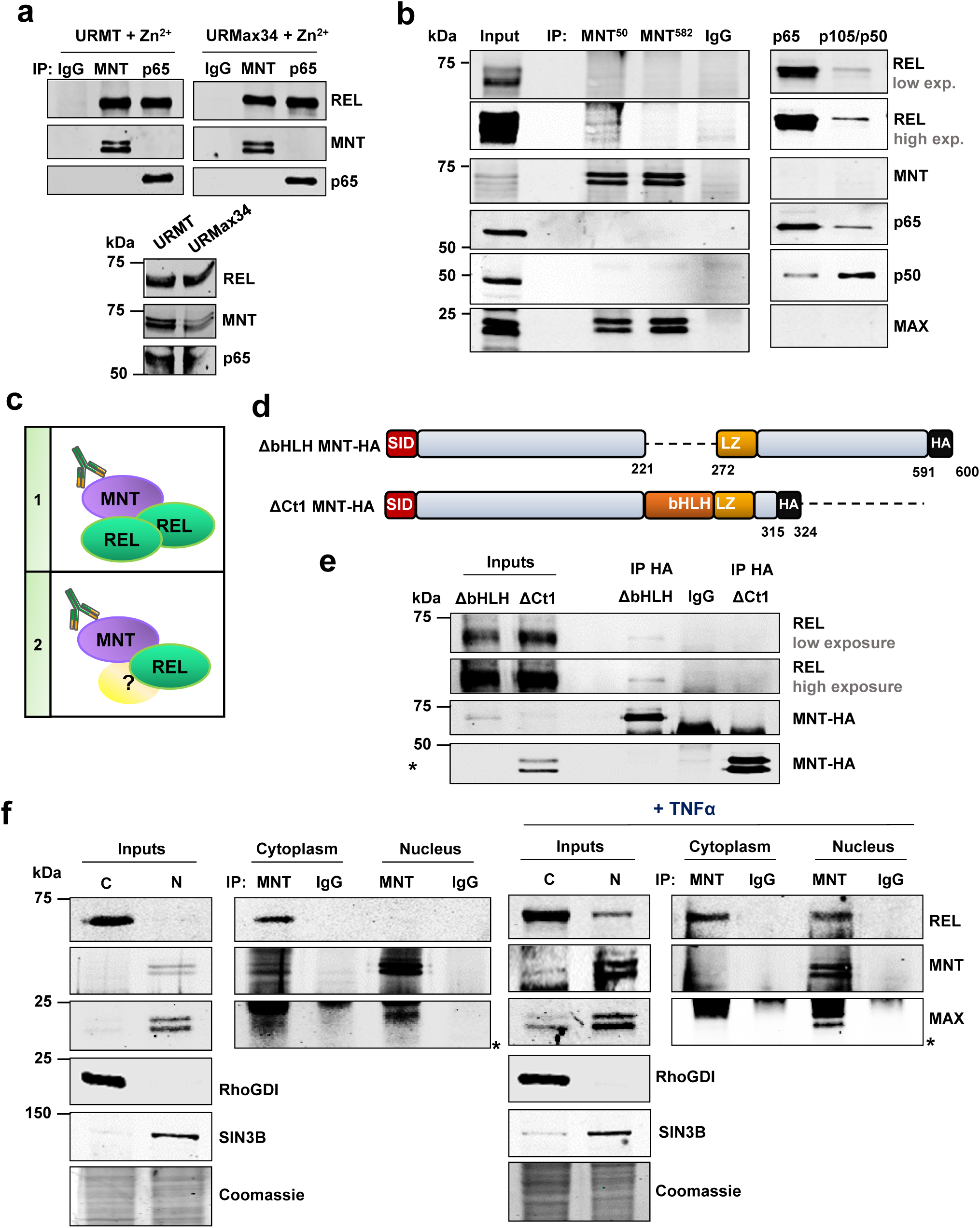
MNT interacts with REL through its C-terminal region in the cytoplasm and the nucleus. **(a)** Co-immunoprecipitation assays with anti-MNT and anti-p65 antibodies (the IgG as a negative control) in URMT and URMax34 after Zn^2+^ treatment. **(b)** Co-immunoprecipitation assay with MNT antibodies (MNT^50^, antibody against the 50 first amino acids of MNT protein; MNT^582^, antibody against the 532-582 amino acids of MNT protein), and with p65 and p105/p50 antibodies, and IgG as a negative control. Two immunoblots for REL are shown (low and high intensity). **(c)** Working hypothesis with two options: (i) MNT bound to REL homodimers, (ii) MNT bound to REL and other yet unknown protein(s). **(d)** Schematic representation of the mouse MNT deletion constructs used for the co-IP assays, ΔbHLH and ΔCt1 MNT-HA. **(e)** C6 cell lysates 48 h after transfection with REL-flag (mouse) and ΔbHLH or ΔCt1 MNT-HA (mouse) were immunoprecipitated with anti-HA antibodies (IgG as negative control). The immunoblot of HA and REL is shown. The asterisk marks the IgG band. **(f)** C6 cells were lysed following the nucleus/cytoplasm fractionation protocol, both in basal conditions or 30 min after treatment with TNFα (100 ng/mL) and immunoprecipitated with an anti-MNT antibody or the IgG (the latter as a negative control). The presence of REL, MNT and MAX was determined in the immunoprecipitates by immunoblot. RhoGDI and SIN3B were analyzed as cytoplasm and nucleus markers, respectively. Coomassie blue was used as a protein loading control for the inputs. Asterisks mark the IgG light chain

### MNT acts as a repressor of the NF-κB’s pathway

We asked if this novel MNT-REL interaction had any impact on the NF-κB signaling. As NF-κB dimers translocate to the nucleus upon the activation of the pathway, we knocked down *MNT* in LoVo cells and performed immunofluorescence assays for REL and p65 to assess their cellular localization. Strikingly, REL accumulated inside the nucleus after *MNT* knockdown, suggesting an activation of the pathway. On the contrary, p65 remained in the cytoplasm regardless of MNT levels (Fig. 4a). This was confirmed by densitometry of the REL and p65 immunofluorescence signals (Supplementary Fig. S3). We next asked whether *MNT* knockdown would cause a release of REL from IκBα by co-IP assays in LoVo cells. The results showed that despite IκBα levels were increased upon *MNT* silencing, its interaction with REL was decreased in the co-IP assay when compared to the control (shScrambled) (Fig. 4b, left). This was confirmed by densitometry (Fig. 4b, right). We also analyzed the protein levels of MNT and NF-κB proteins after *MNT* knockdown. The results showed the increase in p65 and the decrease of REL and p50 protein levels when MNT levels were reduced (Fig. 4c). Since REL was being translocated to the nucleus upon *MNT* knockdown, we monitored the activation of the NF-κB pathway by luciferase assays with a synthetic construct carrying 5 κB binding sites (5x κB-Luc). This is a well-established method to measure NF-κB transcriptional activity^30^. The luciferase activity significantly increased upon MNT knockdown both in LoVo and UR61 cells (Fig. 4d). Our previous work showed that MNT can act not only as MNT-MAX dimers but also as MNT homodimers ^1^. To study whether MNT acts in concert with MAX to repress IκBα promoter, we tested the luciferase activity of the 5x κB-Luc reporter in URMax34 cells, a UR61 derivative in which MAX can be induced by Zn^2+ 1^. MNT knockdown in these cells showed similar elevation of promoter activity in the absence or presence of MAX (Fig. 4d). In addition, we tested the expression of four NF-κB target genes (*BCL-XL/BCL2L1, CCL5, IL6, IL8*) upon MNT knockdown (Fig. 4e, upper panel) and MNT overexpression (Fig. 4e, lower panel). The results confirmed the repressor effect of MNT on the NF-κB pathway.

**Figure 4.**
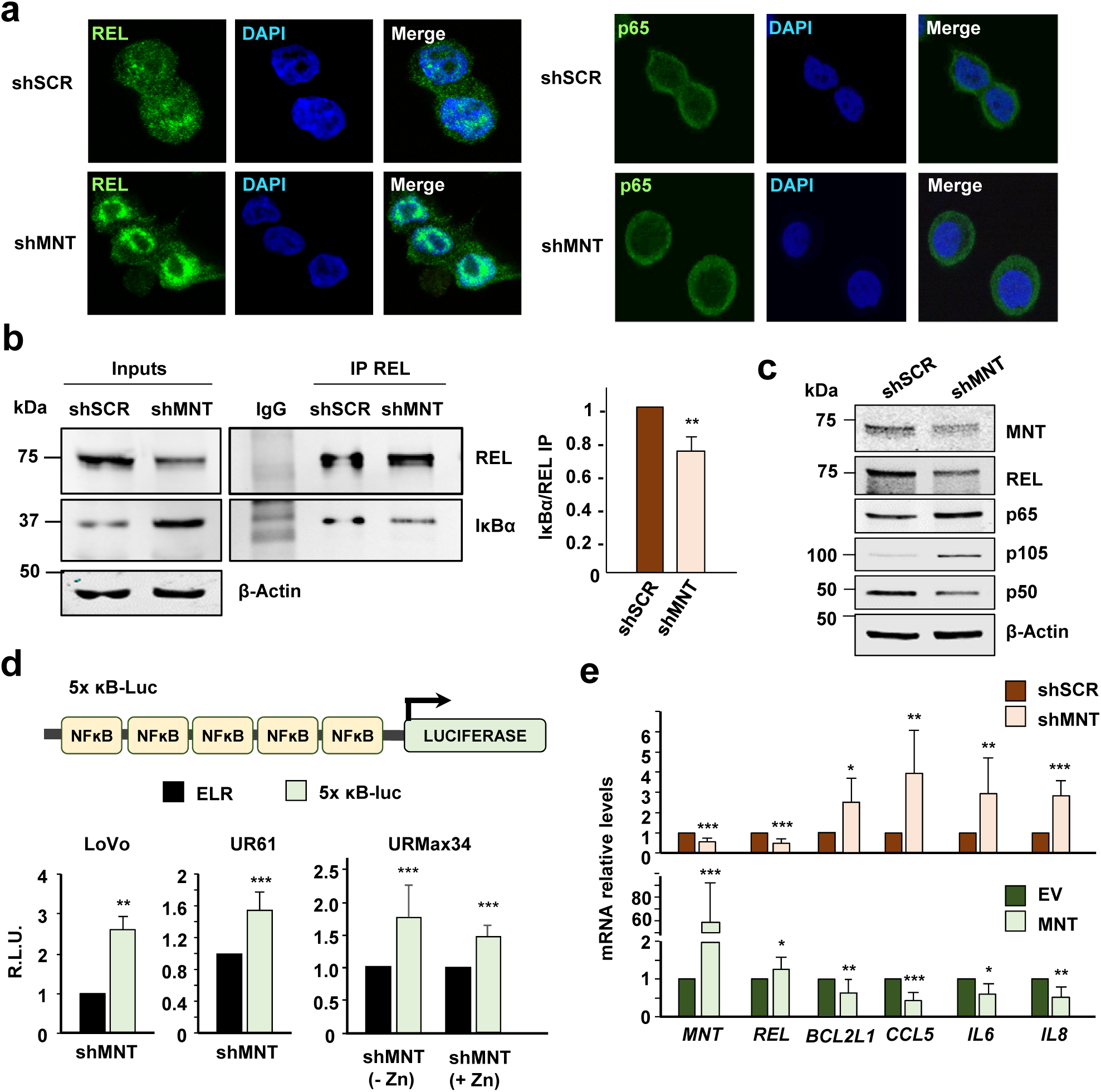
MNT acts as a repressor of the NF-κB’s pathway. **(a)** Immunofluorescence of REL (left panel) or p65 (right panel) in LoVo cells that were infected with lentiviral particles carrying two shRNAs against MNT (shMNT) or a scrambled shRNA (shSCR), and selected with puromycin (1 µg/mL) for 72 h. DAPI as a nuclear marker. **(b)** LoVo cell lysates 72 h after infection with shRNAs against MNT (shMNT) or a scrambled shRNA (shSCR), as a control were immunoprecipitated with anti-REL antibodies (IgG as negative control). The immunoblots of REL and IκBα in the inputs and the IPs are shown. Right graph: the protein levels of the IκBα co-IP versus the REL IP in both conditions were quantified. The data are shown as the mean ± SD, n=2, ** *P* < 0.05. **(c)** Protein levels of MNT, REL, p65, p105/p50 in LoVo cells 72 h after infection with lentiviral particles carrying two shRNAs against MNT or a scrambled shRNA (shSCR), as a control. β-Actin used as a protein loading control. (**d**) Top: Schematic representation of the luciferase reporter driven by five NF-κB binding sites used in these experiments (5xκB-Luc). Bottom: NF-κB-mediated promoter activity. LoVo cells were infected with lentiviral particles carrying two shRNAs against MNT (or a scrambled shRNA as a control, shSCR). Then, 48 h after the infection, cells were selected with puromycin (1 µg/mL) for 72 h and then transfected with the luciferase reporter. Cells were harvested 48 h after the transfection with the luciferase constructs and the luciferasa activity determined. UR61 and URMax34 cells were transfected with the shRNAs and luciferase constructs and harvested 72 h after the transfection for the luciferase assay (URMax34 cells untreated and treated with 100 µM ZnSO_4_ for 24 h). Results are expressed in relative luciferase units (R.L.U.) after normalizing each condition first to the empty luciferase reporter (LER) and then to the shSCR vector. The data are shown as the mean ± SD, n=3 (LoVo), n=4 (UR61 and URMax34). ** *P* <0.05, *** *P* <0.01. **(e)** mRNA levels of *MNT, REL, CCL5, IL6, IL8, BCL2L1* (BCL-XL) in LoVo cells 72 h after infection with shRNAs against MNT (shMNT) or a scrambled shRNA (shSCR) (upper panel) or 48 h after transfection with a MNT overexpressing construct or its corresponding empty vector (EV) (lower panel) relative to RPS14. The data are shown as the mean ± SD, n ≥ 3, * *P* < 0.1, ** *P* < 0.05; *** *P* < 0.01

### MNT directly regulates NFKBIA/IκBα

Given the effects of MNT on the levels of proteins of the NF-κB pathway (Fig. 4c) we asked whether this effect was exerted at the transcriptional level. LoVo cells were transfected with a MNT expression vector to analyze the mRNA levels. The results showed a repression of *RELA* and *NFKBIA* after *MNT* overexpression while *REL* and *NFKB1* expression did not change (Fig. 5a). We focused on *NFKBIA* (IκBα), a repressor of the pathway that is induced by REL^22,31^. To confirm the regulation of *NFKBIA/*IκBα by MNT, we compared *NFKBIA* mRNA levels after overexpressing either wild-type, ΔbHLH (no binding to DNA) or ΔCt1 MNT (unable to interact with REL). The results showed that the repression of *NFKBIA* was not detected with either of the deletion mutants (Fig. 5b). The results suggested a possible regulation of *NFKBIA* by a MNT-REL complex at the transcriptional level. Thus, we studied if MNT was able to repress IκBα through reporter-luciferase assays. Using a construct carrying the promoter of *NFKBIA* (IκBα-Luc), it was found that increased MNT levels led to a repression of *NFKBIA* promoter activity. Consistently with this result, the depletion of MNT through shMNT constructs led to the activation of *NFKBIA* promoter (Fig. 5c).

**Figure 5.**
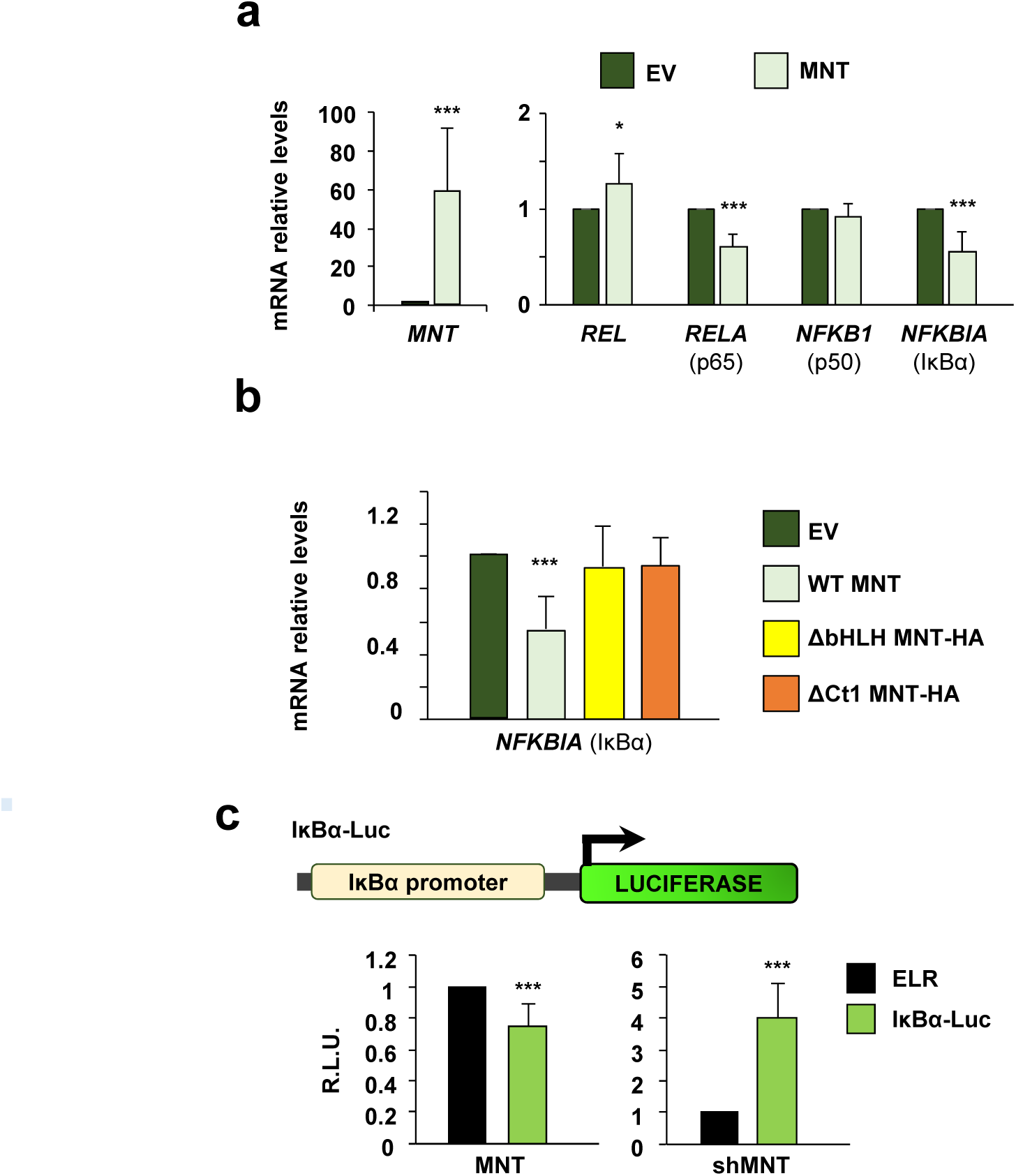
MNT directly regulates *NFKBIA* (IκBα). **(a)** mRNA levels of *MNT, REL, RELA, NFKB1, NFKBIA* in LoVo cells 48 h after transfection with a MNT overexpressing construct or its corresponding empty vector (EV) relative to RPS14. The data are shown as the mean ±SD, n=4, ** *P* < 0.05; *** *P* < 0.01. **(b)** mRNA levels of *NFKBIA* in LoVo cells 48 h after transfection with a WT MNT, ΔbHLH or ΔCt1 MNT-HA overexpressing construct, or their corresponding empty vector (EV) relative to RPS14. The data are shown as the mean ± SD, n=6 (WT), n=3 (mutants) *** *P* < 0.01. **(c)** Top: Schematic representation of the luciferase reporter driven by the human *NFKBIA* (IκBα) promoter (IκBα-Luc) used in this work. For the luciferase assay overexpressing MNT, the luciferase activity was measured 48 h after transfection with the luciferase vectors and MNT expression vector or their corresponding empty vectors (on the left). For the luciferase assay after MNT knockdown, cells were first infected with lentiviral particles carrying two shRNAs against MNT (or a scrambled shRNA as a control, shSCR), selected with puromycin (1 µg/mL) for 72 h and then transfected with the luciferase vectors (on the right). Results are expressed in relative luciferase units (R.L.U.) after normalizing each condition first to the luciferase empty reporter (ELR) and then to the empty vector of MNT (pCMVSport6) (left) or the shSCR (right). The data are shown as the mean ± SD, n = 3 (shMNT), n = 4 (MNT), *** *P* < 0.01.

To test if MNT was bound to the IκBα promoter we analyzed the ChIP-seq data from the ENCODE consortium in the K562 cell line. We observed two coincident peaks of MNT, MYC and MAX on *NFKBIA* gene, suggesting *bona fide* binding sites for dimers of the MYC-MAX-MNT protein family. The transcriptional repressor and MNT partner SIN3A also bound *NFKBIA/*IκBα (Fig. 6a). ChIP experiments for MNT and REL were performed in LoVo cells, which show constitutive NF-κB activation ^32^. We analyzed regions of *NFKBIA* gene −1000 bp to +1000 bp from the TSS, which include some conserved REL-binding sites (Fig. 6b). The results showed MNT and REL binding to *NFKBIA* gene (Fig. 6c). Interestingly, both MNT and REL had a maximum binding at +171/+343 bp, in the first exon of *NFKBIA*. To analyze if both MNT and REL were bound as a complex, we carried out a re-ChIP experiment. For this, we immunoprecipitated chromatin first with anti-MNT and then, MNT-bound chromatin was immunoprecipitated with anti-REL antibodies. The results showed binding of MNT-REL to the +171/+343 region. As a positive control, REL → p50 re-ChIP gave a positive signal on −67/−316, a predicted REL-binding site in that region (Fig. 6d).

**Figure 6.**
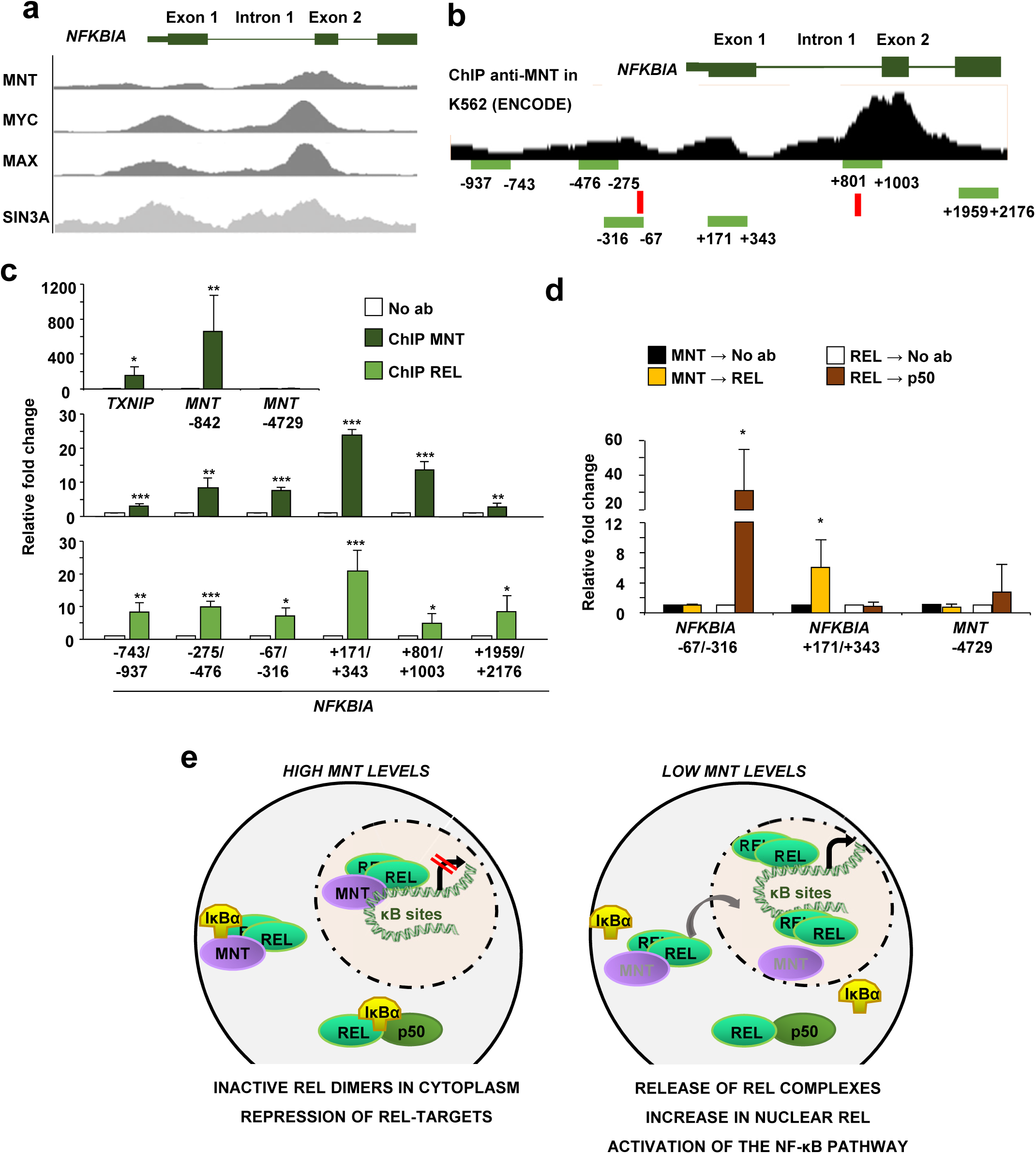
MNT and Rel regulate *NFKBIA*. **(a)** Schematic representation of human *NFKBIA* (IκBα) gene showing the peaks for MNT, MYC, MAX (K562 cell line) and SIN3A (GM78), as published by the ENCODE project consortium (genome-euro.ucsc.edu/). **(b)** Schematic representation of human *NFKBIA* (IκBα) gene showing the peaks for MNT binding in K562 cell line, together with the amplicons analyzed below by ChIP-PCR. The conserved REL-binding sites obtained in the ENCODE project using the TFBS Conserved (tfbsConsSites) track are marked in red rectangles. **(c)** ChIP of MNT (upper panel) or REL (lower panel) in LoVo cells on *NFKBIA* promoter. *TXNIP* and *MNT* −842 used as positive controls and a region upstream MNT promoter (*MNT* −4729), as a negative control for MNT ChIP. The data are shown as the mean ± SD, n=3, ** *P* < 0.05; *** *P* < 0.01 **(d)** Re-ChIP of MNT with REL (MNT → REL) and REL with p50 (REL→p50) in LoVo cells. MNT and REL ChIPs were also re-immunoprecipitated with IgG as a negative control of the technique. The data are shown as the mean ± SD, n ≥ 3, * *P* < 0.1. **(e)** Model of the MNT regulation of NF-κB signaling. Our results indicate that in a high MNT-levels condition, MNT would be possibly regulating NF-κB signaling by retaining REL dimers in the cytoplasm and by forming a complex with REL in the nucleus that would repress the genes that are normally activated by REL dimers. When we silence MNT, REL dimers are released and they translocate into the nucleus, with the consequent increase in NF-κB target genes and the activation of the pathway.

Altogether, the data suggest that MNT inhibits the NF-κB pathway by retaining REL dimers in the cytoplasm and by repressing genes normally activated by REL in the nucleus. When MNT levels drop, NF-κB pathway is activated, as REL dimers are released and they induce the transcription of several NF-κB target genes (Fig. 6e).

## DISCUSSION

Here we report evidences of the involvement of MNT in the regulation of one of the most important signaling pathways, the NF-κB pathway. First, MNT and REL interact in some mouse, rat and human cell lines, and this complex can be found in both cytoplasm and nucleus. Second, *MNT* knockdown triggers REL (but not p65) translocation into the nucleus and the NF-κB pathway activation. Third, MNT-REL complex binds to and regulates a crucial NF-κB target gene: *NFKBIA*/IκBα. The TCGA data showed that concomitant mutations of MNT and REL are rare in tumors, suggesting that they act in the same pathway. Interestingly, MNT and REL interaction is independent of MAX, as it also takes place in MAX-deficient cells and REL was not found in MAX immunoprecipitates. The fact that we did not detect MNT-REL interaction in all the cell lines tested indicates that the interaction may depend on another ancillary protein which expression may vary among cell types. This would affect the efficiency of the co-immunoprecipitation and the Proximity Ligation Assays, leading to a more difficult detection of MNT and REL interaction. Although REL forms heterodimers with p65 or p50 ^22^, we did not detect any of them bound to MNT, which suggests that MNT binds specifically to REL. The C-terminal region of MNT, which is rich in prolines, was necessary for the interaction with REL^3^. These proline-rich regions are usually involved in protein-protein interactions^33^. This agrees with the result showing that REL is found in MNT immunoprecipitates when using an antibody recognizing the first 50 amino acids of MNT but not when using an antibody recognizing the C-terminal domain of MNT. The binding of the antibody to MNT C-terminal domain may disrupt MNT-REL interaction, thus affecting the final result. In addition, we showed that the MNT-REL complex is found in the cytoplasm under basal conditions but also in the nucleus after TNFα stimulation, suggesting a possible transcriptional function of the complex.

*MNT* knockdown provoked the dissociation of the REL-IκBα complexes and the translocation of REL to the nucleus, commonly observed when NF-κB pathway is activated ^34^. In fact, we demonstrate the activation of NF-κB pathway upon MNT silencing, as assessed by a luciferase NF-κB responsive reporter and the increased expression of several NF-κB target genes. In accordance to this result, some NF-κB target genes were down-regulated after *MNT* overexpression, Thus, MNT might be acting as a limiter of NF-κB activity in the absence of specific activators of the pathway.

*NFKBIA* encodes IκBα, which binds to and retains the NF-κB members in the cytoplasm under the absence of stimulatory signals^34^. Once a stimulus is detected by the cell, the NF-κB pathway is activated and IκBα degraded. However, NF-κB also induces a negative feedback loop, leading to the transcription of *NFKBIA*/IκBα. The newly synthesized IκBα enters the nucleus and shuttles NF-κB dimers back to the cytoplasm to terminate transcription^35-37^. Thus IκBα levels analysis is a good method to study the transcriptional activity of NF-κB^38^. ChIP assays show the binding of MNT and REL to the first exon of the IκBα gene and re-ChIP experiments confirm that MNT and REL bind together to that same region. MNT and other MXD proteins exert a transcriptional repressive effect in many genes due to the interaction with SIN3 co-repressor ^2,6^. The ENCODE data shows SIN3A binding to the *NFKBIA/*IκBα gene, which can explain the transcriptional repression of the gene. The fact that MNT can bind and repress IκBα at least in some cell types, opens a new level of regulation of the NF-κB pathway, i.e., the MNT-REL mediated repression of genes otherwise activated by REL.

MNT pro-survival role has been described in several models, although the exact mechanism responsible for that remains unknown^10,39,40^. Here we describe the unexpected discovery of an interaction between MNT and REL, a NF-κB pathway component, which plays a key role in regulating cell homeostasis. It can be hypothesized that MNT would impair REL function by retaining REL dimers in the cytoplasm and also by repressing REL-target genes (Fig. 6e). Thus, the previously described effects of MNT on proliferation could be in part exerted through the regulation of NF-Kb activity. Moreover, it has been described that REL induce MYC expression^41-44^. Considering the MYC-MNT antagonism, it would be possible that MNT would control MYC levels through the inhibition of REL functions. Furthermore, loss of MNT in T-cells leads to a disruption of T-cell development and lymphomagenesis^39,40^. As REL is also important for Th1 cell differentiation^45^, it would be possible that the consequences of MNT on the immune system are related to its interaction with REL

In summary, our results show unprecedented evidence of a MNT-REL interaction, which links the MYC-MNT with the NF-κBB pathways and opens a new path to the understanding of MNT wide functions in cell biology.

## MATERIALS AND METHODS

### Cell culture, transfections and lentiviral transduction

Cell lines were obtained from ATCC and grown in either RPMI-1640 or DMEM (Corning) supplemented with 10 % fetal bovine serum (Gibco, Thermo Fisher Scientific, Waltham, MA, USA), 150 µg/mL of gentamicin and 2 µg/mL of ciprofloxacin. All cells tested negative for Mycoplasma infection by PCR. UR61 derivate from PC12 cells^46^. URMax34 cell line derive form UR61 and express a MAX gene inducible by ZnSO_4_ ^1^, LoVo cells were transfected with ScreenFect A reagent (Screenfect, Eggenstein-Leopoldshafen, Germany), following the manufacture’s indications and using the reagent at 3x µg DNA. C6 were transfected with using PEI reagent (Polysciences, Warrington, PA, USA). UR61 and URMax34 cells were transfected using the Ingenio Electroporation solution (Mirus) in an Amaxa nucleofector. Transfected plasmids were human MNT (pCMVSport6-MNT, Origene Technologies, Rockville, MD, USA); ΔbHLH MNT-HA (murine MNT carrying a deletion of amino acids 221-272 amino acids and tagged with HA) and ΔCt_1_ MNT-HA (murine MNT carrying a deletion of 276 amino acids in the C-terminal region and tagged with HA), both in pcDNA3 with the zeocin resistance gene inserted. Lentiviral production was performed as previously described^47^ and polybrene at 3 µg/mL was used to increase the infection efficiency. Lentiviral particles carried a scrambled short-hairpin RNA as a control, shSCR (SHC016-1EA) or two short-hairpin RNAs against MNT human gene, shMNT (TCR0000234788 and TRCN0000235815), from Sigma-Aldrich, St. Louis, MO, USA. Activation of the NF-κB’s pathway was achieved by TNFα (Peprotech, Rocky Hill, CT, USA) stimulation for 30 min at 25-100 ng/mL.

### RNA extraction and expression analysis

For qPCR, total RNA was isolated using the TRI Reagent® Solution (Invitrogen, Thermo Fisher Scientific, Waltham, MA, USA). The cDNA was generated by reverse transcription (RT) using the iScript (Bio-Rad, Hercules, CA, USA). Quantitative polymerase chain reaction (qPCR) was performed with specific primers (**Supplementary Table 2**) using the iTaq™ Universal SYBR® Green Supermix (Bio-Rad) and CFX ConnectTM Real-Time PCR Detection System (Bio-Rad). RNA was converted into cDNA and analyzed as described ^48^. Levels of mRNA were normalized against RPS14 mRNA levels.

### Immunoprecipitation assays and immunoblot

For the immunoprecipitation assays, lysates were obtained using the 1 % NP-40 IP lysis buffer (50 mM TrisHCl pH 7.5, 150 mM NaCl, 1 % NP-40, 1 mM EDTA pH 8, 0.5 mM EGTA pH 8 and protease and phosphatase inhibitors). For MNT-REL co-IP in LoVo cells, the cells were lysed instead with a mild hypotonic buffer (10 mM HEPES pH 7, 10 mM KCl, 0.25 mM EDTA pH 8, 0.125 mM EGTA pH 8, 0.5 mM spermidin, 0.1 % NP40, 1 mM DTT and phosphatase and protease inhibitors). Total cell lysis, immunoblots and immunoprecipitations (IPs) were performed as described^48^. The antibodies are shown in the **Supplementary Table 3**.

### Preparation of cytoplasmic and nuclear fractions

Cytoplasmic extracts were obtained by a 30 min lysis with a hypotonic buffer (10 mM HEPES pH 7, 10 mM KCl, 0.25 mM EDTA pH 8, 0.125 mM EGTA pH 8, 0.5 mM spermidin, 0.1 % NP40, 1 mM DTT and phosphatase and protease inhibitors). Nuclear extracts were obtained after a 5 min centrifugation at 1500 rpm and lysed with 1 % NP-40 IP lysis buffer (50 mM TrisHCl pH 7.5, 150 mM NaCl, 1 % NP-40, 1 mM EDTA pH 8, 0.5 mM EGTA pH 8 and protease and phosphatase inhibitors). Once the lysates were obtained, the immunoprecipitation and immunoblots were performed as described^48^.

### Proteomic studies

Protein immunoprecipitation was carried out as described in protein immunoprecipitation section except of the elution, which was carried out according to the “On-beads digestion” protocol^49^. Briefly, beads-immunocomplexes were trypsinized, in order to digest the baits and the interacting proteins. After trypsinization, protein samples were purified and finally resuspended in 0.1 % (v:v) trifluoroacetic acid buffer to be analyzed by mass spectrometry on a Q-Exactive mass spectrometer (Thermo Fisher Scientific) connected to an Ultimate Ultra3000 chromatography system (Thermo Fisher Scientific). Mass spectra were analyzed using the MaxQuant Software package of two technical replicates and biological triplicates of the experimental and control samples. Raw data files were searched against a *Rattus norvegicus* (Rat) database (Uniprot RAT), using a mass accuracy of 6 ppm and 0.01 false discovery rate (FDR) at both peptide and protein level.

### Immunofluorescence staining

Adherent cells grown on glass coverslips were fixed with 4 % paraformaldehyde in PBS for 15 min at room temperature. Fixed cells were washed with PBS and permeabilized and blocked with 1 % Triton X-100, 3 % BSA in PBS during 30 min. Then, cells were treated with blocking buffer (3 % BSA; 0.1 % Triton X-100 in PBS) for 20 min, washed with PBS and 0.1 % Triton X-100 in PBS, and incubated overnight at 4°C with the primary antibodies 1:200 diluted in blocking buffer. The slides were incubated for 1 h at room temperature with the secondary antibody conjugated with FITC (Jackson Laboratories, Bar Harbor, ME, USA). The samples were mounted with ProLong Gold Antifade mountant (Thermo Fisher Scientific, Waltham, MA, USA). Confocal images were obtained with a Leica TCS SP5 microscope and processed and quantified using the ImageJ software (https://imagej.nih.gov/ij/download.html). The antibodies used are described in **Supplementary Table 3**.

### *In Situ* Proximity Ligation Assay

In situ Proximity Ligation Assay (PLA) was performed with Duolink *in situ* Red Starter kit Mouse/Rabbit (Sigma-Aldrich, St. Louis, MO, USA) according to manufacturer’s instructions. In situ PLA positive signals were quantified using the ImageJ software. Cell samples were visualized using a Zeiss Axio Imager M1 upright fluorescence microscope. The primary antibodies used are described in **Supplementary Table 3**.

### Luciferase reporters and assays

Cells were transfected with a mix of DNA constructs specific for each experiment. The wild-type human IκBα promoter construct has been previously described ^50^. The firefly luciferase gene reporter vector carrying five NF-κB binding sites was described ^30^. As a control, we used a vector without any specific transcription regulatory sequence. The pRL-null Renilla plasmid (Promega, Madison, WI, USA) was also transfected. Luciferase reporter assays were carried out with the Dual-Luciferase Reporter (DLR) System (Promega), following the manufacturer’s instructions. Luminescence from both luciferase reactions was measured with the Glomax Multi-detection System (Promega). Firefly luminescence values were normalized against Renilla luminescence values. The mean of the duplicates was done, and values were relativized against the empty vector (control). The results were represented as Relative Luciferase Units (R.L.U.).

### Chromatin immunoprecipitation (ChIP and Re-ChIP) assays

Total cell extracts were first lysed with a hypotonic buffer (described in the Nuclear/Cytoplasm Fractionation section) for purifying the nuclear compartment. Then, nuclear lysis and chromatin immunoprecipitation (ChIP) were performed essentially as described ^51^. For the Re-ChIP experiments, we carried out an intermediary step of elution with elution buffer plus 10 mM DTT, 1 h at 37 °C. The resulting DNA was incubated with the second antibody. Immunoprecipitated DNA was purified with the QIAquick PCR Purification Kit (Qiagen, Germantown, MD, USA) and analyzed by qPCR. The SYDH ENCODE project was used as a reference for primer designing on human *NFKBIA* gene (http://genome.ucsc.edu/ENCODE). The primers and antibodies used are described in Supplementary Tables 2 and 3, respectively.

### Survival and mutational analysis

The mutational data shown here was generated using the cBioPortal of Cancer Genomics tool ^28,52^. The survival analysis in cancer prognosis was obtained using the online database OncoLnc (http://www.oncolnc.org/)^53^.

### Statistical analysis

Student’s two-tail t-test was used to evaluate the significance of differences between control and experimental groups. A *P*-value was noted as * *P* < 0.1, ** *P* < 0.05, *** *P* < 0.01.

## Acknowledgments

The work was supported by grant SAF2017-88026-R from Agencia Estatal de Investigación, Spanish Government, to JL and MDD. JL-P and MCL-N were recipients of F.P.U. fellowships from Spanish Government. We are grateful to Jose P Vaqué and Jose L Fernandez-Luna for plasmids, and Rosa Blanco and Sandra Zunzunegui for technical help.

---

**Supplementary figure S1.**
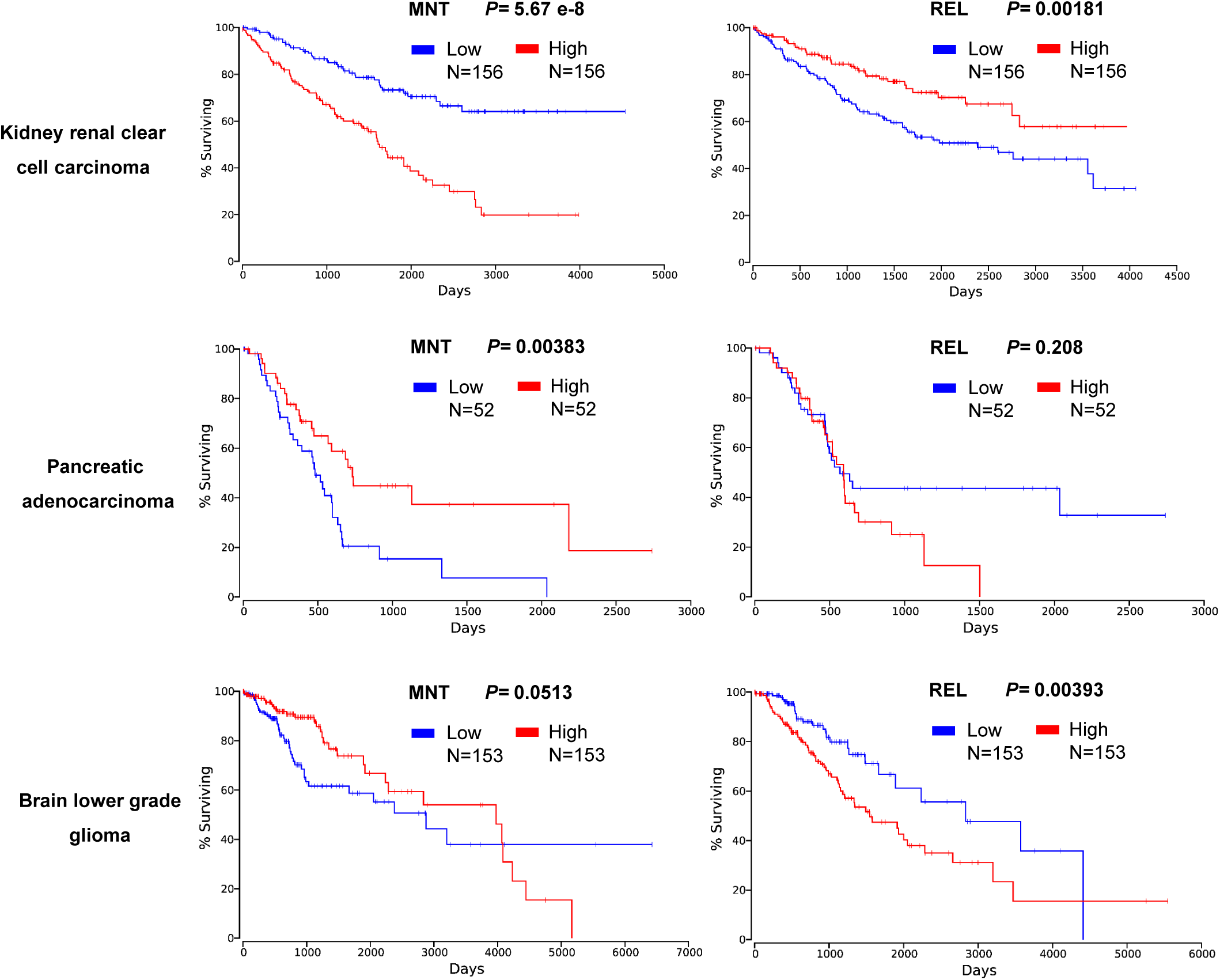
Survival analysis of MNT and REL for kidney renal clear cell carcinoma, pancreatic adenocarcinoma and brain lower grade glioma prognosis using the online database OncoLnc. The number of patients per group and the logrank *P*-value is specified for each plot.

**Supplementary figure S3.**
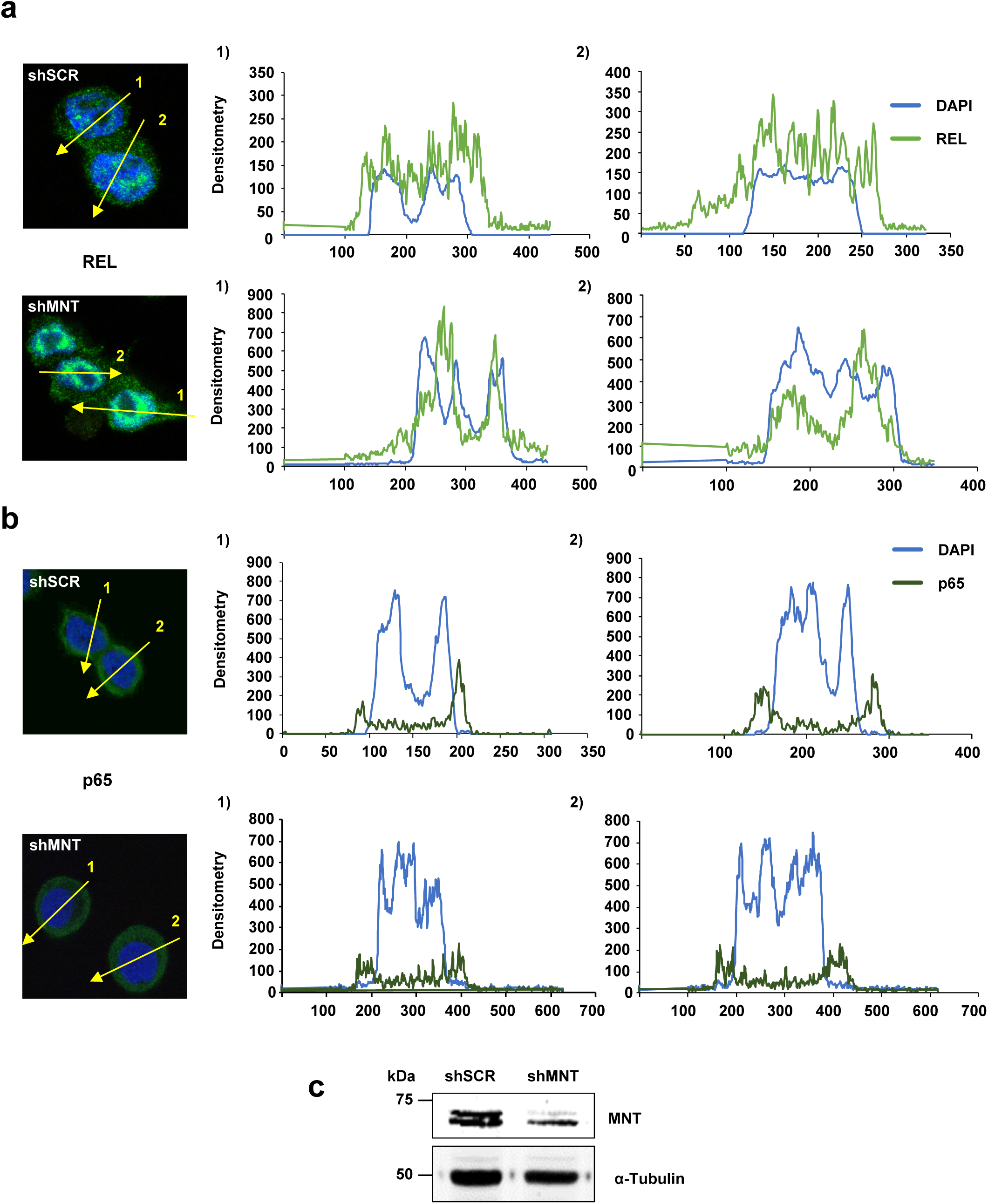
**(a)** REL immunofluorescence and **(b)** p65 immunofluorescence in LoVo cells that were infected with lentiviral particles carrying two shRNAs against MNT (shMNT) or a scrambled shRNA (shSCR) and selected with puromycin (1 µg/mL) for 72 h. The signal was quantified with the ImageJ software. **(c)** Immunoblot of the cells used for the immunofluorescence showing the *MNT* knockdown efficiency. α-Tubulin was determined as a protein loading control.

## References

1. Lafita-Navarro MC, Liano-Pons J, Quintanilla A, et al. The MNT transcription factor autoregulates its expression and supports proliferation in MYC-associated factor X (MAX)-deficient cells. The Journal of biological chemistry 2020; 295: 2001–17.

2. Hurlin PJ, Queva C, Eisenman RN. Mnt: a novel Max-interacting protein and Myc antagonist. Current topics in microbiology and immunology 1997; 224: 115–21.

3. Meroni G, Reymond A, Alcalay M, et al. Rox, a novel bHLHZip protein expressed in quiescent cells that heterodimerizes with Max, binds a non-canonical E box and acts as a transcriptional repressor. The EMBO journal 1997; 16: 2892–906.

4. Meroni G, Cairo S, Merla G, et al. Mlx, a new Max-like bHLHZip family member: the center stage of a novel transcription factors regulatory pathway? Oncogene 2000; 19: 3266–77.

5. Conacci-Sorrell M, McFerrin L, Eisenman RN. An Overview of MYC and Its Interactome. Cold Spring Harbor perspectives in medicine 2014; 4: 1–24.

6. Ayer DE, Lawrence QA, Eisenman RN. Mad-Max transcriptional repression is mediated by ternary complex formation with mammalian homologs of yeast repressor Sin3. Cell 1995; 80: 767–76.

7. Grzenda A, Lomberk G, Zhang JS, Urrutia R. Sin3: master scaffold and transcriptional corepressor. Biochimica et biophysica acta 2009; 1789: 443–50.

8. Yang G, Hurlin PJ. MNT and Emerging Concepts of MNT-MYC Antagonism. Genes (Basel) 2017; 8.

9. Link JM, Hurlin PJ. The activities of MYC, MNT and the MAX-interactome in lymphocyte proliferation and oncogenesis. Biochimica et biophysica acta 2015; 1849: 554–62.

10. Hurlin PJ, Zhou ZQ, Toyo-oka K, et al. Deletion of Mnt leads to disrupted cell cycle control and tumorigenesis. The EMBO journal 2003; 22: 4584–96.

11. Toyo-oka K, Hirotsune S, Gambello MJ, et al. Loss of the Max-interacting protein Mnt in mice results in decreased viability, defective embryonic growth and craniofacial defects: relevance to Miller-Dieker syndrome. Human molecular genetics 2004; 13: 1057–67.

12. Foley KP, McArthur GA, Queva C, Hurlin PJ, Soriano P, Eisenman RN. Targeted disruption of the MYC antagonist MAD1 inhibits cell cycle exit during granulocyte differentiation. The EMBO journal 1998; 17: 774–85.

13. Queva C, McArthur GA, Iritani BM, Eisenman RN. Targeted deletion of the S-phase-specific Myc antagonist Mad3 sensitizes neuronal and lymphoid cells to radiation-induced apoptosis. Molecular and cellular biology 2001; 21: 703–12.

14. Cvekl A, Jr., Zavadil J, Birshtein BK, Grotzer MA, Cvekl A. Analysis of transcripts from 17p13.3 in medulloblastoma suggests ROX/MNT as a potential tumour suppressor gene. Eur J Cancer 2004; 40: 2525–32.

15. Edelmann J, Holzmann K, Miller F, et al. High-resolution genomic profiling of chronic lymphocytic leukemia reveals new recurrent genomic alterations. Blood 2012; 120: 4783–94.

16. Lo Nigro C, Venesio T, Reymond A, et al. The human ROX gene: genomic structure and mutation analysis in human breast tumors. Genomics 1998; 49: 275–82.

17. Vermeer MH, van Doorn R, Dijkman R, et al. Novel and highly recurrent chromosomal alterations in Sezary syndrome. Cancer research 2008; 68: 2689–98.

18. Schaub FX, Dhankani V, Berger AC, et al. Pan-cancer Alterations of the MYC Oncogene and Its Proximal Network across the Cancer Genome Atlas. Cell Syst 2018; 6: 282–300 e2.

19. Nguyen HV, Vandenberg CJ, Ng AP, et al. Development and survival of MYC-driven lymphomas require the MYC antagonist MNT to curb MYC-induced apoptosis. Blood 2020; 135: 1019–31.

20. Burnichon N, Cascon A, Schiavi F, et al. MAX mutations cause hereditary and sporadic pheochromocytoma and paraganglioma. Clinical cancer research : an official journal of the American Association for Cancer Research 2012; 18: 2828–37.

21. Pantaleo MA, Urbini M, Indio V, et al. Genome-Wide Analysis Identifies MEN1 and MAX Mutations and a Neuroendocrine-Like Molecular Heterogeneity in Quadruple WT GIST. Mol Cancer Res 2017; 15: 553–62.

22. Gilmore TD, Gerondakis S. The c-Rel Transcription Factor in Development and Disease. Genes & cancer 2011; 2: 695–711.

23. Zhang Q, Lenardo MJ, Baltimore D. 30 Years of NF-kappaB: A Blossoming of Relevance to Human Pathobiology. Cell 2017; 168: 37–57.

24. Chen IS, Wilhelmsen KC, Temin HM. Structure and expression of c-rel, the cellular homolog to the oncogene of reticuloendotheliosis virus strain T. Journal of virology 1983; 45: 104–13.

25. Gilmore TD, Cormier C, Jean-Jacques J, Gapuzan ME. Malignant transformation of primary chicken spleen cells by human transcription factor c-Rel. Oncogene 2001; 20: 7098–103.

26. Hunter JE, Leslie J, Perkins ND. c-Rel and its many roles in cancer: an old story with new twists. Br J Cancer 2016; 114: 1–6.

27. Hopewell R, Ziff EB. The nerve growth factor-responsive PC12 cell line does not express the Myc dimerization partner Max. Molecular and cellular biology 1995; 15: 3470–8.

28. Cerami E, Gao J, Dogrusoz U, et al. The cBio cancer genomics portal: an open platform for exploring multidimensional cancer genomics data. Cancer Discov 2012; 2: 401–4.

29. Pimentel-Muinos FX, Mazana J, Fresno M. Biphasic control of nuclear factor-kappa B activation by the T cell receptor complex: role of tumor necrosis factor alpha. Eur J Immunol 1995; 25: 179–86.

30. Martin D, Galisteo R, Ji Y, Montaner S, Gutkind JS. An NF-kappaB gene expression signature contributes to Kaposi’s sarcoma virus vGPCR-induced direct and paracrine neoplasia. Oncogene 2008; 27: 1844–52.

31. Chen C, Edelstein LC, Gelinas C. The Rel/NF-kappaB family directly activates expression of the apoptosis inhibitor Bcl-x(L). Molecular and cellular biology 2000; 20: 2687–95.

32. Sakamoto K, Maeda S, Hikiba Y, et al. Constitutive NF-kappaB activation in colorectal carcinoma plays a key role in angiogenesis, promoting tumor growth. Clinical cancer research : an official journal of the American Association for Cancer Research 2009; 15: 2248–58.

33. Kay BK, Williamson MP, Sudol M. The importance of being proline: the interaction of proline-rich motifs in signaling proteins with their cognate domains. FASEB journal : official publication of the Federation of American Societies for Experimental Biology 2000; 14: 231–41.

34. Gilmore TD. Introduction to NF-kappaB: players, pathways, perspectives. Oncogene 2006; 25: 6680–4.

35. Hayden MS, Ghosh S. Shared principles in NF-kappaB signaling. Cell 2008; 132: 344–62.

36. Kanarek N, Ben-Neriah Y. Regulation of NF-kappaB by ubiquitination and degradation of the IkappaBs. Immunol Rev 2012; 246: 77–94.

37. Sun SC, Ganchi PA, Ballard DW, Greene WC. NF-kappa B controls expression of inhibitor I kappa B alpha: evidence for an inducible autoregulatory pathway. Science 1993; 259: 1912–5.

38. Bottero V, Imbert V, Frelin C, Formento JL, Peyron JF. Monitoring NF-kappa B transactivation potential via real-time PCR quantification of I kappa B-alpha gene expression. Mol Diagn 2003; 7: 187–94.

39. Dezfouli S, Bakke A, Huang J, Wynshaw-Boris A, Hurlin PJ. Inflammatory disease and lymphomagenesis caused by deletion of the Myc antagonist Mnt in T cells. Molecular and cellular biology 2006; 26: 2080–92.

40. Link JM, Ota S, Zhou ZQ, Daniel CJ, Sears RC, Hurlin PJ. A critical role for Mnt in Myc-driven T-cell proliferation and oncogenesis. Proceedings of the National Academy of Sciences of the United States of America 2012; 109: 19685–90.

41. Lee H, Arsura M, Wu M, Duyao M, Buckler AJ, Sonenshein GE. Role of Rel-related factors in control of c-myc gene transcription in receptor-mediated apoptosis of the murine B cell WEHI 231 line. The Journal of experimental medicine 1995; 181: 1169–77.

42. Gupta S, Kumar P, Kaur H, et al. Constitutive activation and overexpression of NF-kappaB/c-Rel in conjunction with p50 contribute to aggressive tongue tumorigenesis. Oncotarget 2018; 9: 33011–29.

43. Grumont R, Lock P, Mollinari M, Shannon FM, Moore A, Gerondakis S. The mitogen-induced increase in T cell size involves PKC and NFAT activation of Rel/NF-kappaB-dependent c-myc expression. Immunity 2004; 21: 19–30.

44. Slotta C, Schluter T, Ruiz-Perera LM, et al. CRISPR/Cas9-mediated knockout of c-REL in HeLa cells results in profound defects of the cell cycle. PloS one 2017; 12: e0182373.

45. Visekruna A, Volkov A, Steinhoff U. A key role for NF-kappaB transcription factor c-Rel in T-lymphocyte-differentiation and effector functions. Clin Dev Immunol 2012; 2012: 239368.

46. Guerrero I, Pellicer A, Burstein DE. Dissociation of c-fos from ODC expression and neuronal differentiation in a PC12 subline stably transfected with an inducible N-ras oncogene. Biochemical and biophysical research communications 1988; 150: 1185–92.

47. Lafita-Navarro MC, Blanco R, Mata-Garrido J, et al. MXD1 localizes in the nucleolus, binds UBF and impairs rRNA synthesis. Oncotarget 2016; 7: 69536–48.

48. Garcia-Gutierrez L, Bretones G, Molina E, et al. Myc stimulates cell cycle progression through the activation of Cdk1 and phosphorylation of p27. Scientific reports 2019; 9: 18693.

49. Turriziani B, Garcia-Munoz A, Pilkington R, Raso C, Kolch W, von Kriegsheim A. On-beads digestion in conjunction with data-dependent mass spectrometry: a shortcut to quantitative and dynamic interaction proteomics. Biology (Basel) 2014; 3: 320–32.

50. Algarte M, Kwon H, Genin P, Hiscott J. Identification by in vivo genomic footprinting of a transcriptional switch containing NF-kappaB and Sp1 that regulates the IkappaBalpha promoter. Molecular and cellular biology 1999; 19: 6140–53.

51. Garcia-Sanz P, Quintanilla A, Lafita MC, et al. Sin3b interacts with myc and decreases myc levels. The Journal of biological chemistry 2014; 289: 22221–36.

52. Gao J, Aksoy BA, Dogrusoz U, et al. Integrative analysis of complex cancer genomics and clinical profiles using the cBioPortal. Sci Signal 2013; 6: pl1.

53. Anaya J. OncoLnc: linking TCGA survival data to mRNAs, miRNAs, and lncRNAs. PeerJ Computer Science 2016; 2: e67.

